# Evolutionary tuning of complexin–synaptotagmin fusion clamp governs regulated neurotransmitter release

**DOI:** 10.64898/2026.10.26.761994

**Authors:** Yuanmo Wang, Dipayan Bose, Douglas Griffith, Jeremy S. Dittman, Shyam S. Krishnakumar

## Abstract

Synaptic vesicles must be held in a fusion-ready but restrained state until Ca²⁺ influx triggers release. Synaptotagmin-1 (Syt1) and Complexin (CPX) both contribute to this fusion clamp, yet removing either protein produces opposite effects in different species. Whether this reflects genuine divergence in their molecular roles, or species-specific tuning of a conserved mechanism, has remained unresolved. Here we address this by comparing mammalian and *C. elegans* orthologs in a reconstituted single-vesicle assay that isolates clamping from upstream docking and priming. We find that clamping in both species requires the combined action of CPX and Syt1, with neither sufficient on its own. CPX clamps only with its cognate SNARE complex, a specificity conferred entirely by its accessory helix. The Syt1 orthologs differ markedly in intrinsic clamping strength, which maps to C2B mediated higher-order oligomerization and sets the functional dependence on CPX in each system. These results support a dual-regulator model of exocytosis in which Syt1 and CPX act sequentially and cooperatively through discrete molecular states to control vesicle clamping and Ca²⁺-triggered release. This framework reconciles long-standing functional discrepancies across species and reveals how evolutionary divergence modulates conserved mechanisms governing synaptic fidelity.

## Introduction

The brain functions with extraordinary speed and precision, enabled by rapid and tightly regulated communication between neurons. This communication is mediated by the release of neurotransmitters from synaptic vesicles (SVs) in response to an action potential evoked influx of calcium ions (Ca²⁺) into the presynaptic terminal^1–3^. The molecular machinery underlying neurotransmitter release is well characterized, and its core components are conserved across species^1–4^. SV fusion is catalyzed by the neuronal SNARE (soluble N-ethylmaleimide–sensitive factor attachment protein receptor) proteins^5,6^. As an SV approaches the pre-synaptic plasma membrane, cognate SNARE proteins on the vesicles and target membrane (v- and t-SNAREs, respectively), progressively assemble or ‘zipper’ into a stable complex that provides the energy required for membrane fusion^5,6^.

To ensure the speed and fidelity of neurotransmission, synapses maintain a readily releasable pool (RRP) of docked SVs that are primed to fuse in response to a Ca^2+^ signal^7,8^. Vesicle docking and priming are orchestrated by SNARE associated chaperones, namely Munc13 and Munc18, which guide the SNARE assembly process^4,6,10^. The vesicle associated Ca^2+^-sensor, Synaptotagmin-1 (Syt1) and regulatory factor, Complexin (CPX), have been implicated in ‘clamping’ or reversibly arresting the SNARE zippering at an intermediate stage to maintain SVs in a fusion ready state^2–4^. Maintaining SVs in this restrained yet Ca^2+^-responsive state is essential not only for rapid signaling, but also for preventing synaptic noise and aberrant neurotransmission. However, the respective contribution of Syt1 and CPX in establishing this fusion clamp are not well understood, as disruption of either protein produces strikingly different synaptic phenotypes across species.

In mammalian neurons, Syt1 deficiency potentiates spontaneous fusion while strongly impairing evoked release^11–14^, whereas CPX loss eliminates all modes of vesicular release^15,16^. In *Drosophila*, CPX loss produces a dramatic elevation of spontaneous fusion together with reduced evoked release^17,18^, while Syt1 loss impairs synchronous evoked release and modestly elevates spontaneous release frequency^16–21^. In *C. elegans,* CPX loss likewise elevates spontaneous release while reducing evoked fusion^22–24^, whereas loss of Syt1 reduces both spontaneous and evoked neurotransmitter release^25–27^. These divergent outcomes are difficult to interpret in terms of clamping, as both Syt1 and CPX act at multiple steps of the vesicle cycle, contributing to docking and priming as well as to the fusion clamp^18–20,22–24^, and loss of either protein alters the number of release-ready vesicles alongside their stability.

Investigations into CPX and Syt1 clamping function across species have largely been confined to individual model organisms. To date, no direct cross-species comparisons have been reported for Syt1, and only a limited number exist for CPX^28–30^. Notably, earlier studies, including some from our groups, did not account for the influence of CPX expression levels on their functional outcomes. This is a critical oversight, particularly in light of our recent findings demonstrating that elevated CPX abundance can partially compensate for clamping deficits^31,32^. It therefore remains unresolved whether CPX or Syt1 acts as the primary clamp prior to Ca²⁺ influx, and whether the two proteins have come to operate differently in different species. Resolving this requires a system in which protein identity and abundance are set independently and clamping is measured downstream of docking.

We therefore used a reconstituted single-vesicle fusion assay^32,33^ that meets both requirements. The platform permits direct comparison of orthologous proteins under identical conditions, with rigorous control over rigorous control of all experimental parameters, and tracking the fate of each vesicle from the moment of docking^32–35^. Using this approach, we find that CPX and Syt1 are interdependent components of a single clamping reaction: neither is sufficient alone, and the stability of the clamp reflects their combined contribution. This architecture is conserved across species, whereas the relative contribution of each component is not. In mammals, Syt1 provides the majority of the clamping activity and CPX contributes incrementally; whereas in *C. elegans*, the reduced clamping efficacy of Syt1 renders CPX indispensable. These findings indicate that the divergent requirements for these proteins across species arise from a redistribution of clamping activity rather than from changes in the clamping machinery.

## Results

To directly compare the functional properties of the CPX and Syt1 orthologs, we utilized a biochemically-defined experimental platform capable of tracking individual vesicle docking, clamping, and Ca^2+^-triggered release using confocal fluorescence microscopy^32,33,36^. Building on our previous work^32,33^, the assay conditions was designed as follows: fluorescently labeled (2% ATTO647N-PE ) small unilamellar vesicles bearing v-SNAREs (∼70 copies, unless otherwise noted) ± Syt1 (∼20 copies) were introduced to a suspended lipid bilayer containing pre-assembled t-SNARE complexes (1:400 protein-to-lipid), with CPX (2 µM) added in solution (Figure 1A). Incorporation of the preassembled t-SNARE complex allowed us to bypass the requirement for the SNARE-assembling chaperones Munc18 and Munc13 in this reconstituted system. The fluorescence signal enabled precise tracking of the docking events and fate of individual vesicles. After attaching to the lipid bilayer, vesicles either undocked (indicated by the sudden disappearance of the fluorescence signal) or fused spontaneously (attested by a rapid burst and gradual decrease in fluorescence intensity as the lipids diffuse away), typically within 10 seconds of docking. Alternatively, vesicles transitioned into a stably docked immobile state, classified as ‘clamped’ (Figure 1A). We monitored large ensemble of vesicles (∼200 vesicles per condition) and the ‘docking-to-fusion’ latency time, cumulatively expressed as ‘survival percentage’, provided a direct measure of the strength of the fusion clamp. Because the fate of single docked vesicles were tracked, this analysis allowed examination of the clamping strength, independent of any alteration in the preceding docking process. The functional competence of clamped vesicles was validated by applying 100 µM Ca²⁺ and confirming their ability to undergo Ca^2+^-triggered fusion (Figure 1A). ***CPX and Syt1 act synergistically to clamp SNARE-mediated vesicle fusion across species.*** We first examined the effect of CPX or Syt1, either individually or in combination, on SNARE-mediated vesicle fusion reaction under otherwise identical experimental conditions. In line with our previous findings^32,33^, we found that mammalian CPX1 (mCPX) or mammalian Syt1 (mSyt1) alone were insufficient to clamp neuronal mammalian SNARE (mSNARE)-mediated fusion. Under these conditions, the majority of the docked vesicles either undocked or fused spontaneously, with only ∼5-20% of vesicles remaining stably docked at 10 seconds post-docking (Figure 1B). In contrast, inclusion of both mCPX1 and mSyt1 resulted in ∼90% of vesicles transitioning into a stably ‘clamped’ immobile state (Figure 1B).

**Figure 1.**
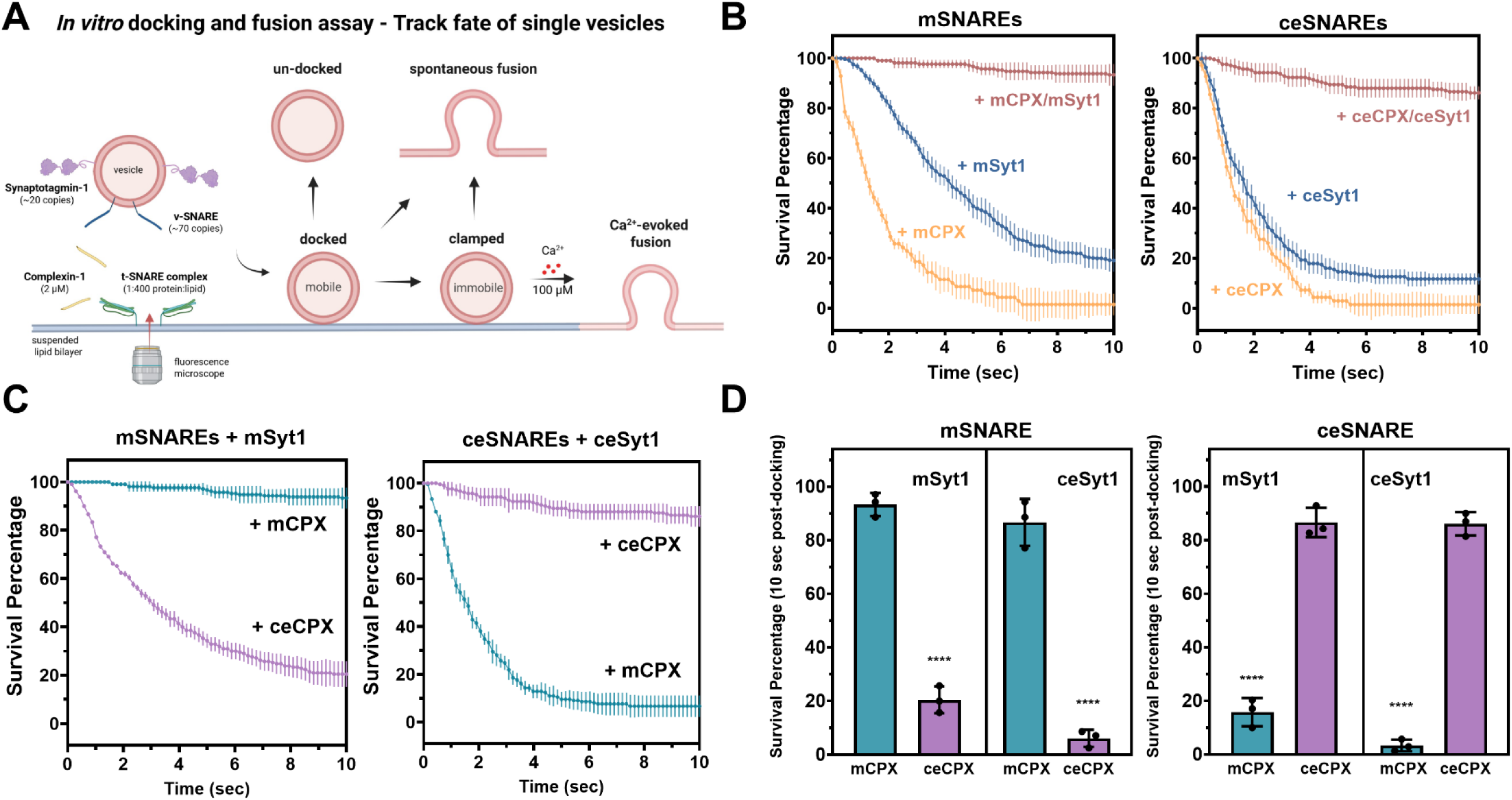
Syt1 and CPX act cooperatively to clamp SNARE-mediated fusion across species. **(A)** Schematic of the in vitro single-vesicle assay. Fluorescently labeled vesicles containing v-SNARE and Syt1 were introduced to a suspended lipid bilayer reconstituted with the t-SNARE complex in the presence of CPX. Docking and the subsequent fate of individual vesicles were monitored by fluorescence microscopy. Docked vesicles either undocked, fused spontaneously, or entered a stably ‘clamped’ immobile state. Clamped vesicles could then be triggered to fuse by Ca²⁺ addition, confirming their functional competence. **(B)** Survival analysis of docking-to-fusion latency shows that CPX and Syt1 were together required to clamp vesicle fusion prior to Ca^2+^ influx in both species, with neither protein sufficient on its own. **(C)** Survival analysis reveals that CPX orthologs clamp efficiently only when paired with their cognate SNARE complex: mCPX clamps mSNAREs but not ceSNAREs, and vice versa, indicating that CPX clamps SNARE-mediated fusion in a species-specific manner. **(D)** End-point analysis (percentage of stably-clamped vesicles at 10 sec post-docking) shows that the con-specific CPX clamping occurs irrespective of the Syt1 isoform used, indicating that selectivity is driven by interactions with the SNARE proteins. Data (mean ± standard deviation) from three independent experiments (N=3; ∼200 vesicles per condition) are shown in all panels. Log-rank (Mantel-Cox) test indicated significant statistical differences (***p<0.001) between survival curves shown in (B) and (C). Two-way ANOVA indicated a statistically significant difference (****p<0.0001) between con-specific and hetero-specific conditions in (D).

A similar requirement for cooperative action was observed for the worm orthologs. In ceSNARE-mediated fusion reactions, only ∼5% of vesicles entered the clamped immobile state in the presence of either worm CPX1 (ceCPX) or worm Syt1 (ceSyt1) alone, whereas ∼90% of docked vesicles remained stably clamped when both ceCPX and ceSyt1 were present (Figure 1B). In all cases, the stably clamped vesicles were functionally competent, with over 90% undergoing rapid fusion upon application of 100 µM Ca²⁺ (Supplementary Figure 1). These results demonstrate that CPX and Syt1 act cooperatively to establish a stable, Ca²⁺-sensitive fusion clamp in both mammalian and invertebrate systems.

### CPX clamping requires Syt1 but is dictated by CPX–SNARE compatibility

Having established that CPX and Syt1 act synergistically to clamp vesicle fusion, we next asked whether this cooperative clamping mechanism is universal or dictated by any species-specific molecular interactions. To address this, we systematically tested clamping efficacy by independently pairing CPX and Syt1 orthologs with SNARE complexes either from the same species (con-specific) or from a different species (hetero-specific), allowing us to dissect how interactions between CPX, Syt1, and SNAREs contribute to the establishment of a fusion clamp.

Survival analysis of docking-to-fusion latency revealed that efficient clamping occurred only when CPX was paired with SNAREs from the same species. Under these conspecific conditions, mCPX robustly clamped vesicle fusion mediated by mSNAREs, and ceCPX showed similarly efficient clamping with ceSNAREs, with ∼90% of vesicles remaining stably docked (Figure 1C). In contrast, heterospecific CPX–SNARE pairings were markedly less effective, with clamping efficiency dropping to ∼10% (Figure 1C). Importantly, this species-specific clamping behavior was observed regardless of which Syt1 ortholog was present (Figure 1D). This suggests that while Syt1 is required to enable CPX-mediated clamping, the species specificity of the clamp is dictated primarily by CPX-SNARE interactions rather than by Syt1 identity. In all conditions where vesicles were stably clamped, subsequent Ca²⁺ addition triggered rapid fusion, confirming that clamped vesicles remained release competent (Supplementary Figure 1). Together, these results demonstrate that clamping of vesicle fusion is not interchangeable across species but instead depends on compatibility between CPX and the underlying SNARE complex.

### CPX accessory helix is necessary and sufficient for its species-specific clamping activity

Because CPX clamping was observed to be species-specific, we sought to identify the structural elements responsible for this divergence between mammalian and worm orthologs. Sequence alignment of the minimal functional domains revealed that the SNARE-binding central helix (CH) is highly conserved (∼80% similarity), whereas the flanking N-terminal accessory helix (AH) and the C-terminal domain (CTD) show substantial divergence (Figure 2A). Both domains have been implicated in CPX function, but their precise roles in clamping remain unclear. To address this, we generated chimeric constructs in which either the AH or CTD was exchanged between mCPX and ceCPX while preserving the remainder of the protein (Figure 2B).

**Figure 2.**
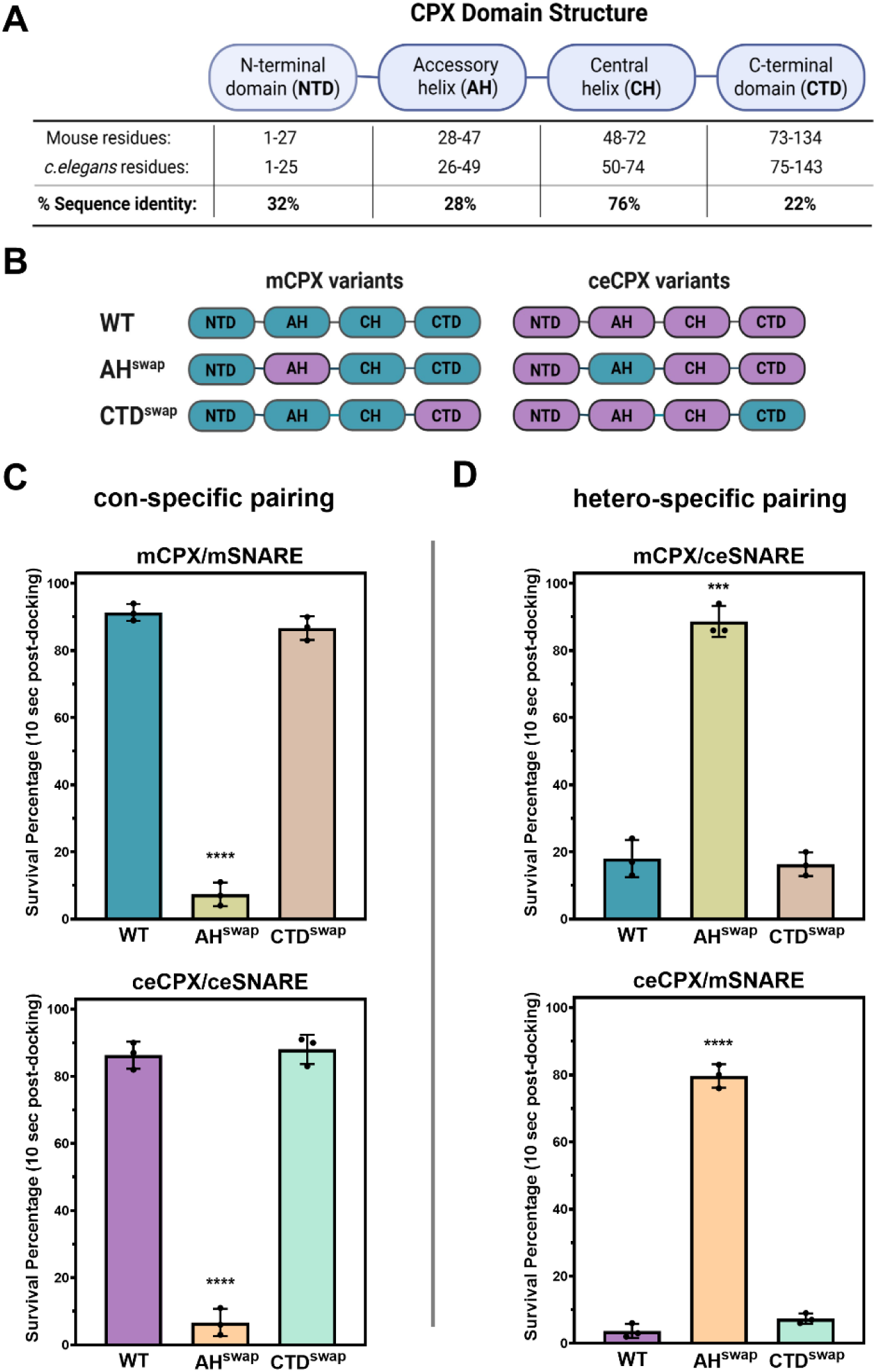
Species-specific interaction of the CPX accessory helix with its cognate SNARE complex underlies clamping selectivity. **(A)** Domain organization of mCPX and ceCPX orthologs . Each contains a highly conserved SNARE-binding motif, the central helix (CH), flanked by a poorly conserved accessory helix (AH) and a C-terminal amphipathic domain (CTD), with all domains contributing to CPX function. (**B**) Chimeric mCPX and ceCPX proteins were generated by swapping either the AH (AH^swap^) or CTD domain (CTD^swap^), leaving the remainder of the protein unchanged. **(C)** End-point analysis (percentage of stably clamped vesicles at 10 s post-docking) shows that swapping the AH domain, but not the CTD domain abolishes CPX-mediated clamping with species-specific SNARE pairing. (**D**) Notably, the AH^swap^ alone was sufficient to confer clamping function under hetero-specific conditions . These results indicate that the AH domain is both necessary and sufficient to determine the species-specific clamping function of CPX orthologs. Data (mean ± standard deviation) from three independent experiments (N=3; ∼200 vesicles per condition) are shown. Two-way ANOVA indicated a statistically significant difference between the AH^swap^ condition and the WT or CTD^swap^ conditions (****p<0.0001; ***p<0.001).

Functional assays showed that swapping the AH (AH^swap^) abolished clamping activity even under conspecific conditions, reducing the fraction of clamped vesicles by >80%. In contrast, CTD swaps (CTD^swap^) had little effect on clamping efficiency, indicating that the CTD does not substantially contribute to species-specific function (Figure 2C). Strikingly, swapping the AH domain also conferred clamping under heterospecific conditions – while wild-type and CTD^swap^ proteins exhibited poor clamping (∼10–20%) when paired with non-cognate SNAREs, AH^swap^ chimeras exhibited robust clamping activity, with ∼90% of vesicles remaining docked and unfused (Figure 2D). In all cases, clamped vesicles were fusion-competent, undergoing rapid Ca²⁺-triggered release (Supplementary Figure 2). Control experiments confirmed that AH or CTD domain swaps did not affect protein folding or SNARE-binding affinity relative to wild-type CPX (Supplementary Figure 3). Taken together with previous studies^17,22,32,37^, these findings indicate that the accessory helix is necessary for CPX clamping and sufficient to determine its species specificity. Clamping therefore depends on compatibility between AH and the cognate SNARE complex, raising the possibility that the two have co-evolved to preserve this compatibility.

### Divergent clamping efficacy of Syt1 orthologs modulates CPX function across species

Together, our findings demonstrate that while the clamping activity of CPX is species-specific, the underlying clamping mechanism is highly conserved. How, then, can we reconcile the strikingly different synaptic phenotypes observed upon CPX or Syt1 loss in mammalian versus invertebrate neurons? To address this, we quantified the intrinsic clamping activity of CPX and Syt1 orthologs separately under conspecific conditions, by varying the number of v-SNARE molecules per vesicle (∼70, ∼35, or ∼18 copies) while holding constant either CPX concentration (2 µM) in the absence of Syt1, or Syt1 density (∼20 copies per vesicle) in the absence of CPX.

Strikingly, neither mCPX nor ceCPX were able to clamp vesicle fusion in the absence of Syt1, as the majority of docked vesicles under all conditions either undocked or underwent spontaneous fusion (Figure 3A, 3B). The failure of CPX to clamp fusion across SNARE stoichiometries and ortholog systems indicates that stable fusion clamp formation cannot be achieved by CPX alone. In contrast, Syt1 orthologs exhibited an intrinsic ability to clamp SNARE-mediated fusion, albeit with markedly different efficacy. As v-SNARE copy number decreased, mSyt1 exhibited progressively stronger clamping activity, stabilizing ∼25%, ∼50%, and ∼85% of vesicles at the respective densities tested (Figure 3C). In comparison, ceSyt1 was substantially less effective under identical conditions, with only ∼1%, ∼12%, and ∼50% of vesicles remaining stably docked (Figure 3D). This indicates that mammalian Syt1 is a more potent fusion clamp compared to the worm ortholog. Importantly, this difference was observed with both mammalian and worm SNAREs (Supplementary Figure 4), indicating that it reflects intrinsic molecular properties of the Syt1 orthologs rather than differences in SNARE identity.

**Figure 3.**
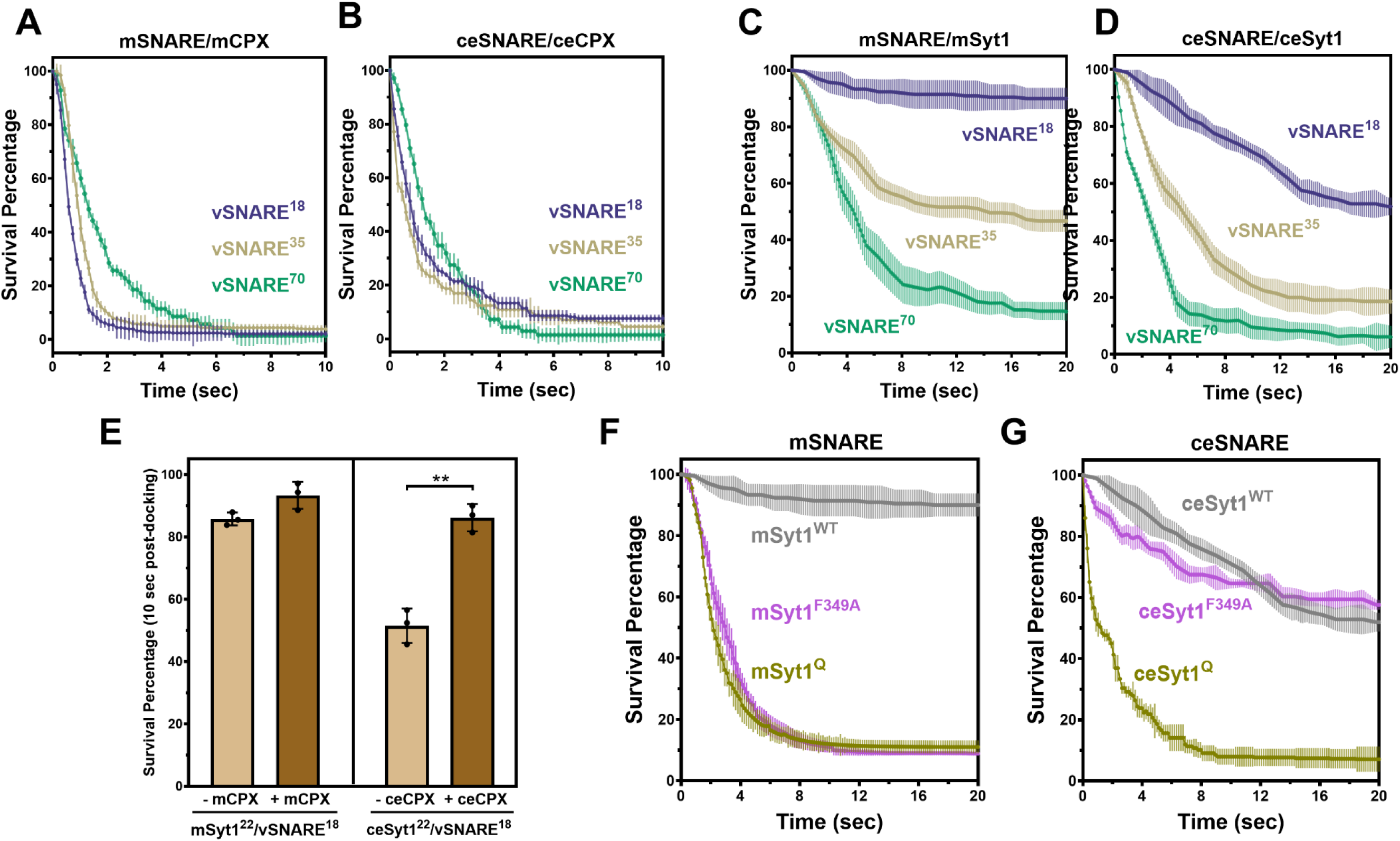
Divergent C2B-mediated oligomerization shapes the clamping ability of Syt1 orthologs and sets the CPX requirement. The intrinsic clamping capacity of CPX and Syt1 orthologs was assessed by systematically varying the v-SNARE copy number per vesicle (∼70, ∼35, or ∼18 copies) while maintaining a constant Syt1 density (∼20 copies per vesicle). (**A,B**) Survival analysis of docking-to-fusion latency under con-specific SNARE conditions reveals that, in the absence of Syt1, neither mCPX (**A**) nor ceCPX (**B**) can clamp vesicle fusion on its own. (**C, D**) Similar analysis in the absence of CPX reveals that both Syt1 orthologs clamp SNARE-mediated fusion intrinsically, as vSNARE copy number decreases but with markedly different efficacy. Notably, mSyt1 (**C**) acts as a more potent fusion clamp compared to ceSyt1 (**D**) under all conditions tested. **(E)** End-point analysis measuring the percentage of stably clamped vesicles at 10 s post-docking under low v-SNARE (∼18 copies/vesicle) condition, shows that the addition of mCPX has minimal effect on the number of ‘clamped’ vesicles relative to mSyt1 alone. In contrast, the inclusion of ceCPX significantly increases the fraction of stably docked vesicles. These results indicate that the CPX functional requirements are tuned by the Syt1 orthologs across species. (**F, G)** Effect of disrupting SNARE ‘primary interface’ binding or oligomerization of C2B domain on clamping by Syt1 orthologs under low v-SNARE conditions in the absence of CPX. For mammalian Syt1 (**F**), both the primary-interface mutant (mSyt1^Q^; R281A, E265A, Y338W, R368A, R366A) and the oligomerization-destabilizing substitution (mSyt1^F346A^) strongly impair clamping, with ∼60% of docked vesicles undocking or fusing spontaneously. For worm Syt1 **(G)**, the corresponding primary-interface mutant (ceSyt1^Q^; R301A, E315A, Y358W, R418A, R416A) eliminates clamping, whereas the oligomerization mutant (ceSyt1^F366A^) is indistinguishable from wild-type. Together, these results identify C2B-mediated oligomerization as the molecular feature distinguishing the clamping efficacy of the two Syt1 orthologs. Data (mean ± standard deviation) from three independent experiments (N=3; ∼200 vesicles per condition) are shown for all panels. Log-rank (Mantel-Cox) test indicated no significant differences between curves in (A) and (B), and significant differences (***p<0.001) between curves in (C), (D), (F) and (G). Welch’s t-test comparing conditions without and with CPX also showed significant difference (** p<0.01) in (E).

The functional contribution of Syt1 and CPX to establish the fusion clamp were strongly dependent on the number of available SNARE complexes and diverged most clearly under the conditions most likely to apply *in vivo*. Although the precise SNARE stoichiometry required for SV exocytosis remains uncertain, converging evidence suggests that a limited number of SNARE complexes assemble beneath docked vesicles and are sufficient to drive Ca²⁺-evoked synchronous neurotransmitter release^38–40^. Under such low SNARE conditions (∼18 copies of v-SNARE per vesicle), mSyt1 alone clamped ∼85% of docked vesicles, with mCPX providing only a modest enhancement (∼93%). In contrast, ceSyt1 alone clamped ∼50% of vesicles, and the addition of ceCPX increased the clamped fraction to ∼85% (Figure 3E). The requirement for CPX in establishing a stable clamp is therefore set by the clamping efficacy of the Syt1 ortholog present, providing a potential basis for the divergent synaptic phenotypes observed across species.

Notably, the Ca²⁺-evoked fusion probability of vesicles clamped by mSyt1 alone was lower than vesicles clamped by mSyt1 together with mCPX, and this difference was most pronounced at low and physiologically-relevant Ca²⁺ concentrations (Supplementary Figure 5). This indicates that beyond its role as a fusion clamp, CPX also enhances the efficacy of Syt1 mediated Ca²⁺-triggered exocytosis. The equivalent comparison could not be made for the worm system, as the high rate of spontaneous fusion under ceSyt1-only conditions precluded reliable measurement of Ca²⁺-evoked fusion probability.

#### Molecular determinants of divergent Syt1 clamping efficacy across species

We next sought to delineate the molecular properties underlying the differential clamping efficacy of Syt1 orthologs. Previous structural and functional studies of mammalian Syt1 have shown that interaction of the C2B domain with the SNARE complex through the ‘primary’ binding interface ^33,41,42^ and C2B-mediated oligomerization^14,43^ both contribute to formation of the fusion clamp. Consistent with this, selective disruption of Syt1-SNARE interaction at the primary interface (R281A, E295A, Y338W, R368A, R366A; mSyt1^Q^)^41,42^ abolished clamping, with only ∼5% of docked vesicles remaining stably clamped under low SNARE conditions (∼18 copies of v-SNARE per vesicle) in the absence of CPX (Figure 3F). Destabilization of mSyt1 oligomerization by a single point mutation (F346A)^43^ similarly impaired clamping, with ∼ 15 % of docked vesicles remaining clamped under identical conditions (Figure 3F).

The C2B domains of mammalian and C. elegans Syt1 share a high degree (∼85%) of sequence similarity and notably all residues implicated in primary SNARE binding and oligomerization are conserved. We therefore generated and tested the corresponding mutations in ceSyt1 under identical conditions. Disruption of the primary SNARE binding interface (R301A, E315A, Y358W, R418A, R419A; ceSyt1^Q^)^27^ eliminated the ceSyt1 fusion clamp, indicating that SNARE engagement is required for ceSyt1 mediated clamping (Figure 3G). In contrast, mutation of the conserved oligomerization residue (F369A) had no detectable effect on clamping efficiency (Figure 3G). Together, these results suggest that the two orthologs share a conserved mode of SNARE engagement, and that their divergent clamping efficacy is likely due to a differing capacity for C2B-mediated higher-order oligomerization, which strengthens the fusion clamp.

## Discussion

Our systematic comparison of mammalian and invertebrate CPX and Syt1 orthologs under biochemically defined conditions reveals a conserved clamping mechanism, built from the cooperative action of these two proteins, together with species-specific differences in how each contributes to it. Critically, the assay measures clamping downstream of docking, as each vesicle is tracked from the moment it docks, so clamp stability is assessed independently of the preceding steps. This matters because both Syt1 and CPX act at multiple stages of exocytosis, from docking and priming through to clamping and Ca²⁺-triggered release. The reconstituted system therefore provides a direct measure of the intrinsic clamping efficacy of each protein and each ortholog.

Our data shows that CPX requires Syt1 to clamp SNARE-mediated membrane fusion and prevent premature vesicle release in both species. The clamping function relies on interactions between the CPX accessory helix and partially assembled SNARE complexes, a mechanistic feature that appears conserved across evolution. Despite this shared core principle, both components are tuned to their species context. CPX is evolutionarily matched to its cognate neuronal SNARE proteins, with sequence divergence in the accessory helix conferring species-specific selectivity. Syt1 orthologs differ in intrinsic clamping efficacy, which in turn sets the extent to which CPX is required for establishing the fusion clamp.

### Species-specific tuning of CPX–SNARE compatibility

Our chimera experiments identify the accessory helix (AH) as the determinant of CPX–SNARE compatibility, with the c-terminal domain (CTD) making no detectable contribution to species specificity. Previous studies have established that deletion of the AH abolishes CPX clamping^17,22,32,37^ and our current data extend this by showing that the AH is also sufficient to determine species specificity. The ability of AH swaps to restore robust clamping in heterospecific environments is particularly notable, indicating that the AH serves as the critical ‘compatibility module’ between CPX and its SNARE machinery. This highlights how subtle sequence divergence can fine-tune regulatory proteins to their native fusion apparatus and suggests that co-adaptation of CPX and SNARE proteins may represent an evolutionary mechanism that preserves the fidelity of neurotransmitter release.

Previous structural and biochemical studies have provided key insights into the AH–SNARE interactions essential for the clamping function of mCPX^37,44–46^. Under prefusion conditions, the mCPX AH domain engages both SNAP25 and VAMP2 simultaneously, with SNAP25 binding mediated primarily by hydrophobic contacts and VAMP2 binding driven by electrostatic interactions^45,46^. Whether ceCPX employs a similar mechanism remains unclear. Owing to the high sequence divergence between mammalian and worm AH domains, conventional sequence alignment is uninformative. Helical wheel analysis, however, indicates that two features of the mammalian helix are preserved in the worm AH, namely a hydrophobic patch corresponding to the mammalian hydrophobic face, and a high content of acidic residues, although the basic residues (e.g., Arg/Lys) proposed to contribute to v-SNARE binding are not apparent (Supplementary Figure 6). Whether one or both of these conserved features underlies AH-mediated clamping, and whether the worm AH engages its cognate SNAREs through the same surfaces or by an alternative mechanism, will require further structural and biochemical analysis to resolve.

The CTD is an important contributor to CPX function ^23,31^ but does not confer species specificity. Rather, the CTD enhances clamping efficiency by promoting CPX localization to vesicles, thereby increasing its effective concentration and facilitating access to assembling SNARE complexes^31,32^. This activity is retained in our system, as loss of the CTD raises the CPX concentration required for effective clamping^32^, indicating that the assay remains sensitive to CTD function. In our reconstituted system, however, CPX is present free in solution, and the minimal composition of the assay makes CTD-dependent positioning less critical than it would be in the cellular milieu. The absence of an effect of CTD swaps on species specificity therefore reflects the domain’s distinct role rather than an insensitivity of the assay. In this way, the AH defines compatibility with the cognate SNARE machinery, while the CTD supports effective engagement of CPX with the vesicle fusion apparatus.

### Divergent clamping efficacy of Syt1 orthologs

A second source of species specificity lies in Syt1 itself. The two orthologs differ substantially in intrinsic clamping strength, with mammalian Syt1 a considerably more potent fusion clamp than its worm counterpart. In mammals, the robust mSyt1 clamp reduces the requirement for mCPX in suppressing spontaneous fusion, consistent with the limited effect of CPX loss on spontaneous release in mammalian synapses. In worms, the weaker ceSyt1 clamp renders CPX indispensable for this purpose, accounting for the pronounced elevation of spontaneous fusion in *cpx*-null animals. Syt1 thus sets the functional framework within which CPX operates.

Our mutational analysis points to C2B-mediated oligomerization as the molecular basis of this difference: the two orthologs share a requirement for primary-interface SNARE engagement, but only mammalian Syt1 clamping is sensitive to disruption of higher-order assembly. This divergence is unexpected, given the overall similarity of their C2B domains. We mutated the residue known to mediate higher-order assembly in mSyt1^14,43^, together with the corresponding residue in ceSyt1, and that the same substitution has different consequences in the two proteins is consistent with ceSyt1 not accessing the higher-order states this residue stabilizes, implying that factors beyond this contact determine whether such assembly occurs. Careful structural and biochemical analysis comparing mSyt1 and ceSyt1 will be required to determine the key factors involved. Nonetheless, the capacity for higher-order oligomerization appears to have diverged evolutionarily between the two orthologs, providing a means of strengthening the fusion clamp without altering the SNARE-binding interface they share.

### A dual-regulator model of exocytosis

Our results further show that CPX not only reinforces the Syt1-mediated clamp but also enhances Ca²⁺-triggered exocytosis of these vesicles. This activating role of CPX is well established, as it has been shown to promote release in both mammalian and invertebrate synapses, acting on spontaneous as well as evoked fusion^16,17,22,47^. This indicates that CPX and Syt1 act cooperatively, with Syt1 serving as the Ca²⁺ sensor that establishes a regulated fusion state and CPX facilitating efficient release upon Ca²⁺ influx. Such a dual role would allow CPX to optimize synaptic transmission by maintaining low spontaneous activity while promoting rapid, synchronous release in response to physiological Ca²⁺ signals.

Building on these findings, we propose a ‘dual-regulator model’ of exocytosis in which Syt1 and CPX act cooperatively and sequentially to guide vesicles through distinct molecular states that balance fusion readiness with restraint (Figure 4). Syt1 engages the release machinery first: it is recruited to the vesicle docking site through PI(4,5)P₂ binding ^48,46^ and can associate with SNAP-25 even prior to ternary SNARE assembly^50^. As zippering proceeds, Syt1 engages the partially assembled complex to establish an ‘*intermediate*’ state [*IS*], in which vesicles are morphologically docked and partially clamped. These vesicles remain susceptible to premature spontaneous release but are competent for Ca²⁺-triggered exocytosis.

**Figure 4.**
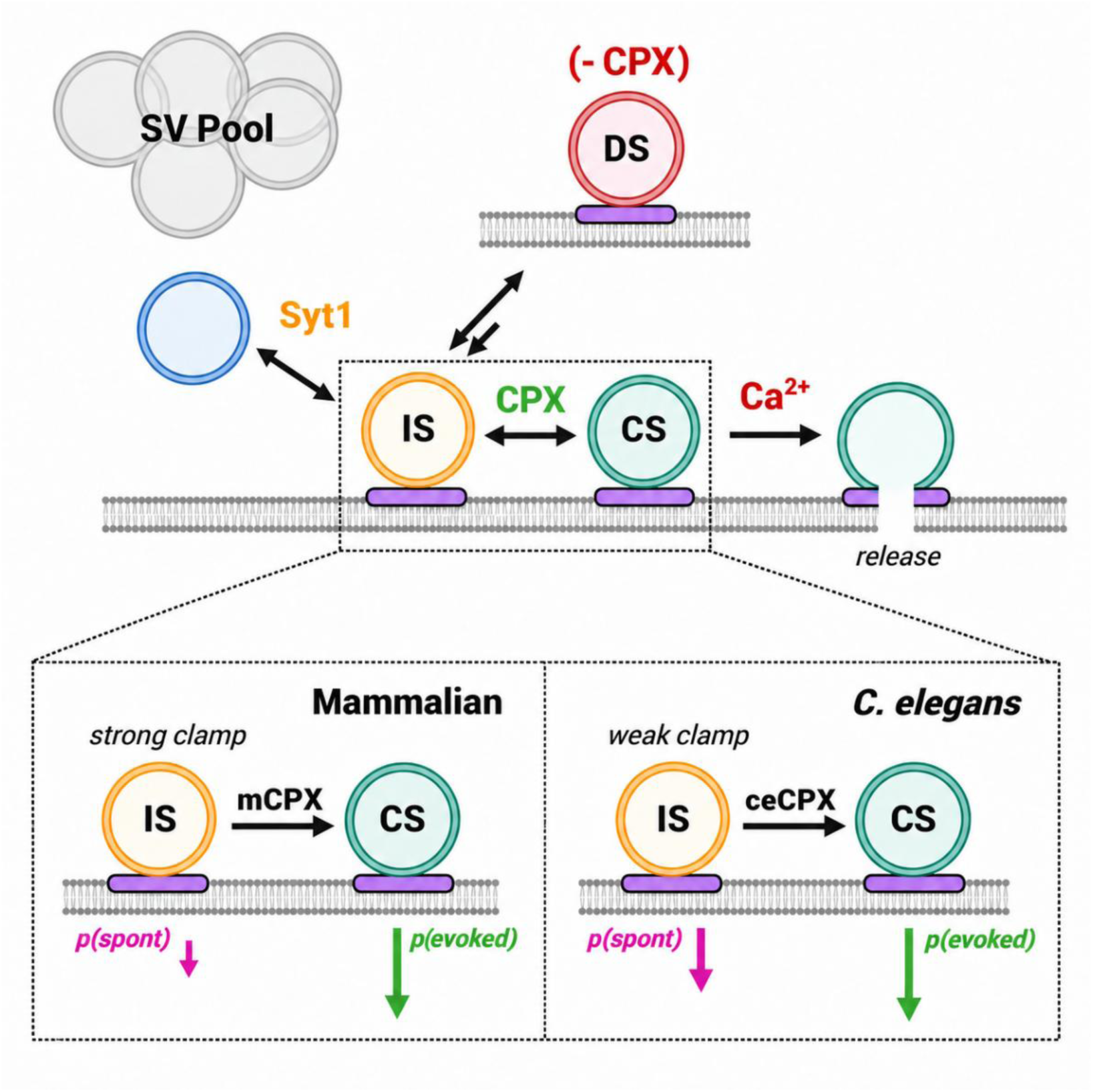
Dual-regulator model of exocytosis. Syt1 and CPX act cooperatively and sequentially at distinct stages of the fusion pathway to coordinate vesicle docking, clamping and Ca²⁺-triggered release. Syt1 engages partially zippered SNARE complexes to establish an intermediate state (IS), in which vesicles are docked and partially clamped, remaining Ca²⁺-responsive but prone to spontaneous fusion. CPX binding converts the IS into a clamped state (CS) by stabilizing the partially zippered SNARE complex, producing a release-ready but fusion-inhibited state. Ca²⁺ influx releases the clamp and drives rapid, synchronous fusion. In the absence of CPX, vesicles in the IS can instead drift into a dysfunctional state (DS) in which the SNARE complex is misconfigured and fusion competence is lost. Inset shows the species-specific tuning of IS-to-CS transition. The strong mammalian Syt1 clamp stabilizes vesicles within the IS and limits spontaneous fusion, so mCPX contributes only incrementally. The weaker ceSyt1 clamp leaves vesicles prone to spontaneous fusion, rendering ceCPX essential for establishing a stable clamp. Spontaneous and evoked fusion probabilities are indicated by magenta and green arrows, respectively, with arrow length reflecting relative probability.

CPX can associate only at this stage, since its central helix binds the interface between syntaxin and VAMP2, which forms as the two begin to zipper^45,51^. CPX binding then drives the transition from the IS to a ‘*clamped*’ state [*CS*]. By stabilizing the partially zippered SNARE complex through its accessory helix, CPX both locks vesicles into a release-ready but fusion-inhibited state and preserves the machinery in a state competent to complete zippering when the clamp is released. Upon Ca²⁺ influx, Syt1 rapidly releases the fusion clamp, allowing SNAREs to complete zippering and drive membrane fusion resulting in neurotransmitter release. A key feature of this model is that the stability of the [*IS*], is not fixed, as it is set by the intrinsic clamping strength of the Syt1 ortholog. The relative contributions of the two regulators are thus quantitatively tunable. This accounts for the divergent phenotypes observed across species when either protein is lost.

In the absence of CPX, vesicles accumulate in the [*IS*], but are not consolidated into the [*CS*], and a fraction of SNAREs may instead transition into ‘*dysfunctional*’ states [*DS*], in which the release machinery is misconfigured, and fusion efficacy is reduced (Figure 4). This does not alter the number of morphologically docked vesicles but lowers their fusion probability except under strong stimulation, such as elevated Ca²⁺ or hypertonic sucrose. CPX loss therefore disrupts the synchrony of evoked release and reduces neurotransmission fidelity in both species, consistent with its role in facilitating Ca²⁺-triggered fusion described above. The consequences for spontaneous release, by contrast, are strongly species-dependent, reflecting the balance between two effects: unclamping, which promotes spontaneous fusion, and [*DS*], accumulation, which reduces the pool competent to fuse by any route. In mammalian synapses, the robust mSyt1 clamp limits the first effect, so the second predominates and all modes of release are reduced. In *C. elegans*, the weaker ceSyt1 clamp leaves vesicles largely unrestrained, and unclamping outweighs [*DS*], accumulation, producing marked increases in spontaneous fusion (Figure 4). Species-specific differences in the [*IS* → *DS*] transition rate may further contribute to these divergent outcomes.

Loss of Syt1 produces a distinct phenotype, preventing vesicles from entering the [*IS*] at all. Because Syt1 also contributes to docking and priming, this reduces the number of release-ready vesicles in addition to removing the clamp, and the resulting phenotype reflects both effects. In *C. elegans* synapses, where Syt1 has been implicated in vesicle priming and little functional redundancy exists^25,26^, the dominant consequence is a reduction in releasable vesicles and hence in both spontaneous and evoked release. In mammalian synapses, Synaptotagmin-7 partially compensates by maintaining near-normal numbers of docked vesicles^52^. So, in the absence of Syt1, these vesicles are unclamped and fuse spontaneously, since CPX alone cannot establish a stable clamp. The model therefore accommodates both outcomes: whether Syt1 loss elevates or lowers spontaneous release depends on how many vesicles remain available to fuse.

This ‘dual-regulator’ model provides a unified framework in which the coordinated actions of Syt1 and CPX establish two sequential checkpoints, offering a flexible regulatory architecture that allows synapses to tune release probability and spontaneous activity through differential modulation of either component. Although the present assay employs preassembled t-SNARE complexes, the roles of Syt1 and CPX described here are reproduced in a fully reconstituted system including Munc13 and Munc18^34^. Nonetheless, physiological studies directly comparing the clamping efficacy of mammalian and worm Syt1 will be needed to test the model further.

Overall, this work addresses a long-standing puzzle of why loss of either CPX or Syt1 produces such divergent synaptic phenotypes across species. Our data suggest that in both cases the answer is not that the clamping machinery differs between species, but that its two components have been tuned to their species context in distinct ways. This underscores the importance of considering evolutionary context and molecular co-adaptation between interacting partners when dissecting mechanisms of synaptic transmission and highlights the value of cross-species approaches in uncovering the organizational principles that govern conserved neural functions.

## Resource availability

All data supporting the findings of this study are available within the manuscript and the supplemental information. This study does not generate original code. Any additional information required is available on request to the lead contact. All materials generated in this study will be available upon request.

## Acknowledgements

This work was supported by NIH grant (R01NS143927) to J.S.D s S.S.K and International Postdoc Fellowship (2021-06643) from Swedish Research Council to Y.W.

## Author contributions

S.S.K and J.S.D conceived the study; Y.W. and D.B. carried out in vitro functional analysis; D.G. contributed unique reagents and carried out biochemical analysis; Y.W., D.B., and S.S.K analyzed the data; S.S.K wrote the paper, with inputs from Y.W. and J.S.D. All authors read and reviewed the manuscript.

## Methods

### Materials

We utilized the following previously described ^32,33^ clones: full-length mammalian VAMP2 (human His^6^-SUMO-VAMP2, residues 1–116), full-length mammalian t-SNARE complex (mammalian His^6^-SNAP25B, residues 1–206 and rat Syntaxin1A, residues 1–288), CPX orthologs (human His^6^-Complexin 1, residues 1–134; C. elegans His^6^-Complexin, residues 1–143), wild type mammalian Syt1 (rat Synaptotagmin1-His^6^, residues 57-421) and SNARE non-binding (R281A, E295A, Y338W, R368A, R366A; mSyt1^Q^) and oligomerization (F349A, Syt1F^346A^) mutants. In addition, we generated and used the following clones: full length C. elegans t-SNAREs complex (His^6^-ric-4a, residues 1–208 and unc-64, residues 1–261), full length *C. elegans* v-SNARE (His^6^-Sumo SNB1, residues 1-106), wild type *C. elegans* Synaptotagmin-1 (His^6^-Snt1; residues 1-441 and oligomerization mutant (ceSyt1^F366A^). We synthesized the chimeric mCPX/ceCPX proteins (AH^swap^ and CTD^swap^) as well as SNARE-nonbinding ceSyt1 (R301A, E315A, Y358W, R418A, R419A; ceSyt1^Q^) in the above-described background from Azenta/Genewiz Inc. (South Plainfield, NJ). The lipids used in the study: 1,2-dioleoyl -snglycero-3-phosphocholine (DOPC), 1,2-dioleoyl-snglycero-3-(phospho-L-serine) (DOPS), and phosphatidylinositol 4, 5-bisphosphate (PIP2) were purchased from Avanti Polar Lipids (Alabaster, AL). ATTO647N-DOPE and ATTO465-DOPE were obtained from ATTO-TEC, GmbH (Siegen, Germany). All other research materials and consumables, unless otherwise specified, were sourced from Sigma-Aldrich (St Louis, MO) and Thermo Fisher Scientific (Waltham, MA)

### Expression and Purification of Proteins

All proteins were expressed and purified using a bacterial expression system, as previously described^32,33^. Briefly, transformed E. coli BL21(DE3) cells (Novagen, Madison, WI) were cultured in Luria broth medium and induced with 0.25 mM IPTG (isopropyl β-D-1-thiogalactopyranoside) overnight to drive protein expression. Cells were harvested by centrifugation, and pellets expressing soluble CPX proteins (WT, chimeras, and mutants) were lysed directly using a cell disruptor (Avestin, Ottawa, Canada) in ‘lysis’ buffer containing 50 mM HEPES, pH 7.4, 400 mM KCl, 4% Triton X-100, Benzonase (5 µL/2 L culture), 10% glycerol, 1 mM dithiothreitol (DTT), and EDTA-free Complete protease inhibitor cocktail (1 tablet/50 mL; Merck, Rahway, NJ).

For transmembrane proteins (e.g., SNAREs and Syt1), pellets were resuspended in membrane isolation buffer containing 50 mM HEPES, pH 7.4, 400 mM KCl, 1 mM DTT, EDTA-free protease inhibitor cocktail (1 tablet/50 mL), and 0.1 mg/mL lysozyme. After disruption using the cell disruptor, the homogenate was centrifuged at 1,000 rpm for 10 min to remove debris. The supernatant containing the membrane fraction was ultra-centrifuged at 100,000 × g for 30 min (45Ti rotor, Beckman Coulter, Atlanta, GA). The resulting membrane pellets were resuspended in ‘lysis’ buffer and shaken for 2 h to solubilize proteins.

All lysates were clarified by ultra-centrifugation (100,000 × g, 30 min, 45Ti rotor), and supernatants were incubated overnight with pre-equilibrated Ni–NTA resin. For soluble CPX proteins, the resin was washed with ‘wash’ buffer containing 25 mM HEPES, pH 7.4, 150 mM KCl, 1 mM DTT, and 25 mM imidazole. For SNARE proteins, wash buffer was supplemented with 1% (w/v) n-octyl-β-D-glucopyranoside (OG). For Syt1 orthologs, the resin was first washed with wash buffer supplemented with 1% Triton X-100 and 20 mM CaCl₂, followed by a second wash with wash buffer containing 1% Triton X-100. The Syt1-bound resin was treated with Benzonase (Millipore Sigma, Burlington, MA) for 1 h at room temperature, then washed with ‘wash’ buffer containing 1% Triton X-100 to remove nucleotide contamination.

Elution was performed as follows: CPX proteins were cleaved from the Ni–NTA resin using thrombin in ‘elution’ buffer (25 mM HEPES, pH 7.4, 400 mM KCl, 0.2 mM TCEP, 10% glycerol). t-SNAREs, worm v-SNARE, and Syt1 orthologs were eluted with 500 mM imidazole in elution buffer supplemented with 1% OG (t-SNARE, worm v-SNARE) or 0.1% Triton X-100 and 1 mM MgCl₂ (Syt1 orthologs). Mammalian VAMP2 was cleaved from the resin using SUMO protease in the ‘elution’ buffer containing 1% OG.

CPX proteins were further purified using a HiLoad 75 gel filtration column, while Syt1 proteins were subjected to ion-exchange chromatography (Mono S column) in elution buffer. Protein concentrations were determined using the Bradford assay (Bio-Rad, Hercules, CA) with bovine serum albumin as the standard. Purity was verified by SDS–PAGE. All proteins were flash-frozen and stored at −80 °C in 10% glycerol.

### Vesicle Preparation

Small unilamellar vesicles including v-SNARE and Syt1 were prepared using a rapid detergent dilution and dialysis method, followed by purification via discontinuous Optiprep gradient ultracentrifugation, as previously described^32,33^. Note: Vesicles containing worm v-SNAREs were treated with SUMO protease to remove the SUMO tag prior to the final rehydration step. The lipid composition, designed to mimic synaptic vesicles, consisted of 88% DOPC, 10% DOPS, and 2% ATTO647N-PE. In most experiments, protein-to-lipid input ratios were 1:100 for v-SNAREs and 1:250 for Syt1. Based on previously characterized reconstitution efficiency and protein orientation ^33^, each vesicle was estimated to contain ∼70 outward-facing copies of v-SNARE and ∼20 outward-facing copies of Syt1, approximating native synaptic vesicle density^53^. To generate vesicles with varied v-SNARE copy numbers, v-SNARE protein-to-lipid input ratios were adjusted to 1:200 or 1:400 yielding vesicles containing ∼35 or ∼18 v-SNARE copies, respectively.

### Suspended Lipid Bilayer Formation

Giant unilamellar vesicles (GUVs) containing t-SNAREs were generated using the osmotic shock protocol^32,33,36^. The GUV lipid composition, mimicking the presynaptic plasma membrane, comprised 80% DOPC, 15% DOPS, 3% PIP2, and 2% ATTO465-PE. The t-SNARE complex was reconstituted at a protein-to-lipid ratio of 1:200, yielding a final concentration of 1:400. GUVs were ruptured on freshly plasma-cleaned Si/SiO₂ chips patterned with a regular array of 5 µm diameter holes in HEPES buffer (25 mM HEPES, 125 mM KCl, 1 mM DTT, 5 mM MgCl₂, pH 7.4). Bilayers were extensively washed with the same buffer containing 1 mM MgCl₂. Bilayer fluidity was confirmed in each experiment by FRAP of ATTO465 fluorescence.

### Single-Vesicle Fusion Assays

Docking and fusion experiments were performed as previously described^32,33^. Vesicles (∼100 nM lipid) and CPX (2 µM final) were added to the suspended bilayer and incubated for 5 min. ATTO647N-PE fluorescence was used to track individual vesicles, quantifying docking, post-docking diffusion, docking-to-fusion delays, and spontaneous fusion. Docked, immobile vesicles remaining un-fused during the observation period were classified as “clamped.” Excess vesicles were removed by buffer exchange, and 100 µM CaCl₂ was added to measure Ca²⁺-triggered fusion of pre-docked vesicles. Movies were acquired at 147 ms/frame, capturing ∼70 vesicles per experiment, at 37°C on an inverted laser scanning confocal microscope (Leica SP5) equipped with a multi-wavelength argon laser (488 nm), diode lasers (532 and 641 nm), and a long-working-distance 40× water-immersion objective (NA 1.1). Emission was spectrally separated and collected using photomultiplier tubes.

### CPX-SNARE Binding Assay

The t-SNARE complex (10 µM) were covalently coupled to NHS-activated-Sepharose beads (Thermo Fisher Scientific, Waltham, MA) following manufacturer’s instruction and extensively washed using HEPES buffer (25 mM HEPES, pH 7.4, 140 mM KCl, 0.2 mM TCEP, 10% glycerol) to remove unbound protein. The t-SNARE-coupled beads were then incubated with 10 µM CPX (WT or chimera variants) in the presence of 60 µM v-SNARE overnight at 4°C. After incubation, the beads were thoroughly washed with HEPES buffer, and bound proteins (CPX s v-SNAREs) were eluted using the SDS loading buffer, followed by boiling at 95°C. The samples were analyzed by SDS-PAGE and visualized by Coomassie blue staining.

