## Supplementary Information for "Evolutionary tuning of complexin–synaptotagmin fusion clamp governs regulated neurotransmitter release"

<sup>1</sup>Nanobiology Institute, Yale University, West Haven, CT, USA; <sup>2</sup>Department of Neurology, School of Medicine, Yale University, New Haven, CT, USA; <sup>3</sup>Department of Psychiatry and Neurochemistry, Institute of Neuroscience & Physiology, The Sahlgrenska Academy at the University of Gothenburg, Mölndal, Sweden; <sup>4</sup>Department of Biochemistry and Biophysics, Weill Cornell Medical College, New York, NY, USA.

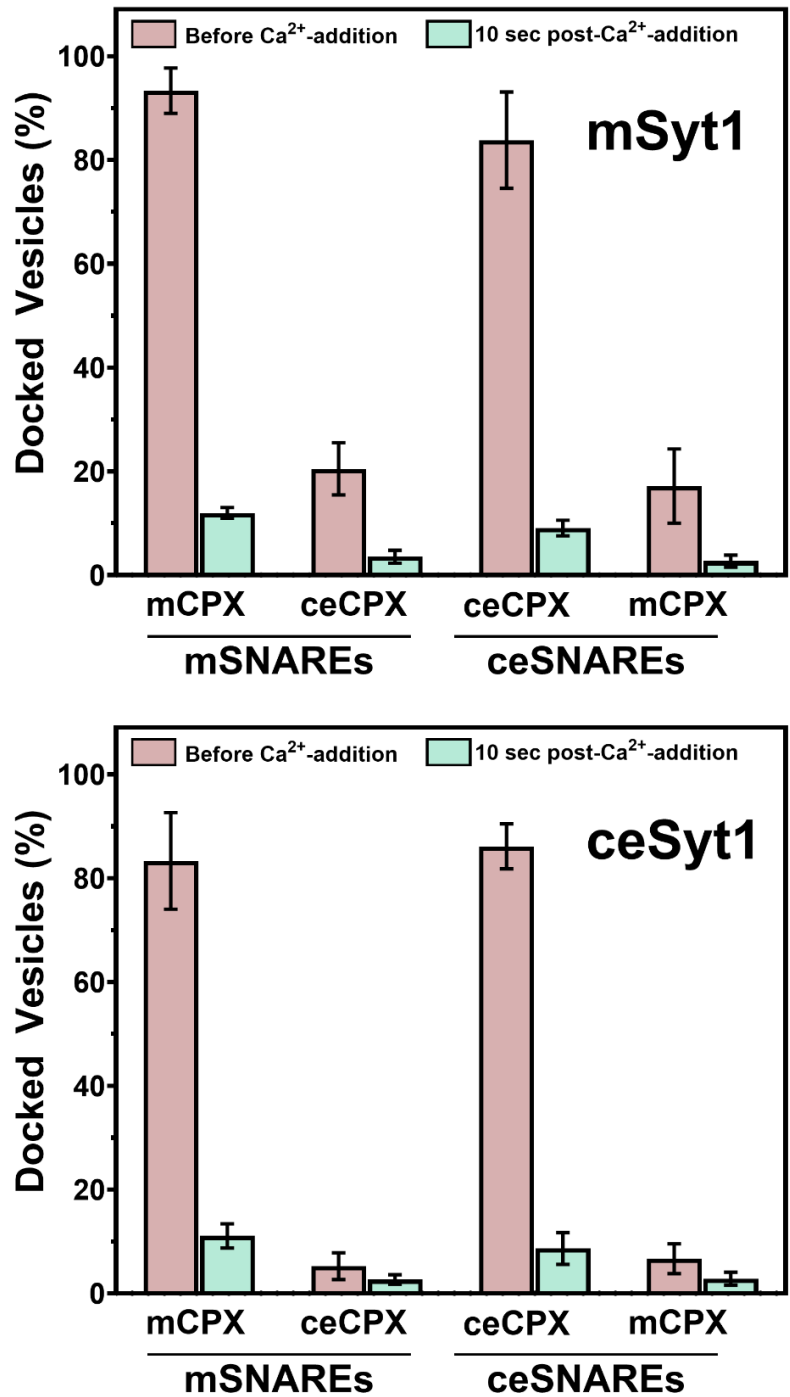

**Supplementary Figure 1.** Fusion analysis demonstrates that the majority of ‘clamped’ vesicles under both con-specific and hetero-specific CPX/SNARE conditions (Figure 1) remain functionally competent, as they can be triggered to fuse upon addition of 100 $\mu\text{M}$ $\text{Ca}^{2+}$. The fraction of docked immobile vesicles is shown before $\text{Ca}^{2+}$ addition (red bars) and 10 seconds after $\text{Ca}^{2+}$ addition (green bars) for both mSyt1 (top) and ceSyt1 (bottom). Data represent mean ± SD from three independent experiments (~200 vesicles per condition).

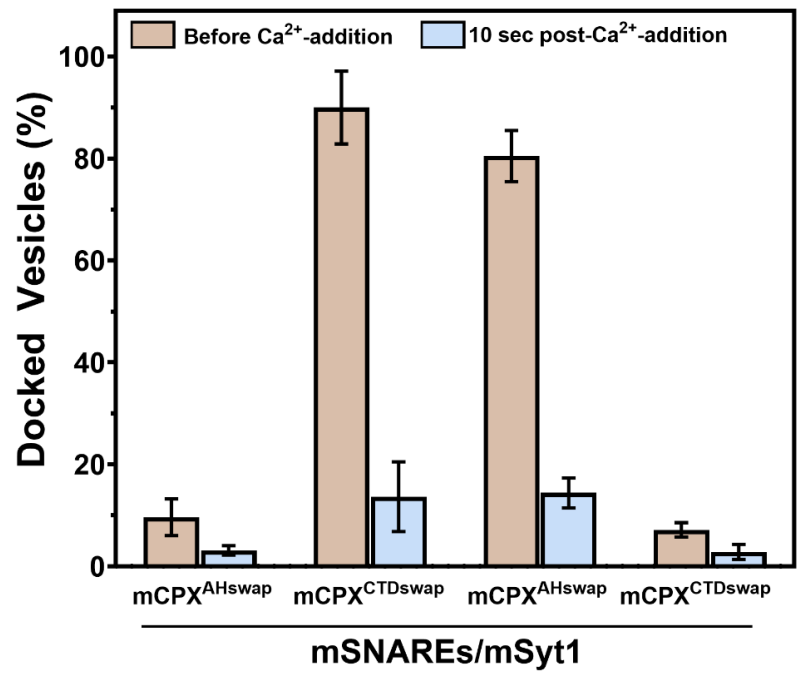

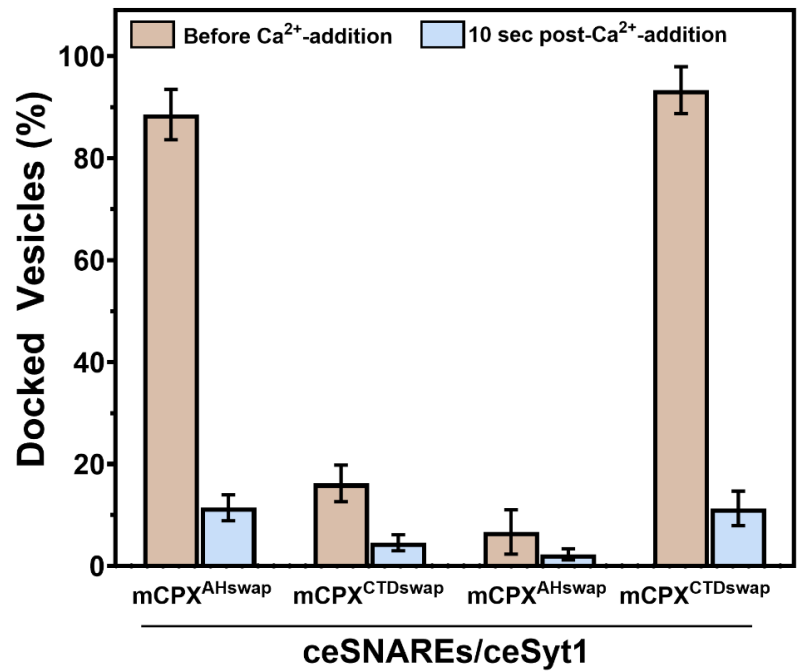

**Supplementary Figure 2.** Fusion analysis indicates that vesicles clamped by AH/CTD domain-swapped mCPX and ceCPX variants, under both conspecific and heterospecific conditions (Figure 2), retain Ca<sup>2+</sup> responsiveness, with 100 μM Ca<sup>2+</sup> triggering fusion of the majority of docked vesicles. Data are shown as the fraction of docked vesicles before Ca<sup>2+</sup> addition (brown bars) and 10 seconds after addition (blue bars), presented as mean ± SD from three independent experiments (~200 vesicles per condition).

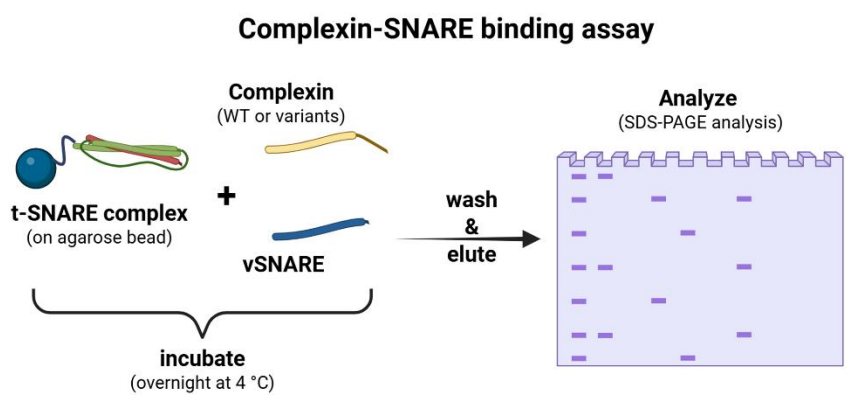

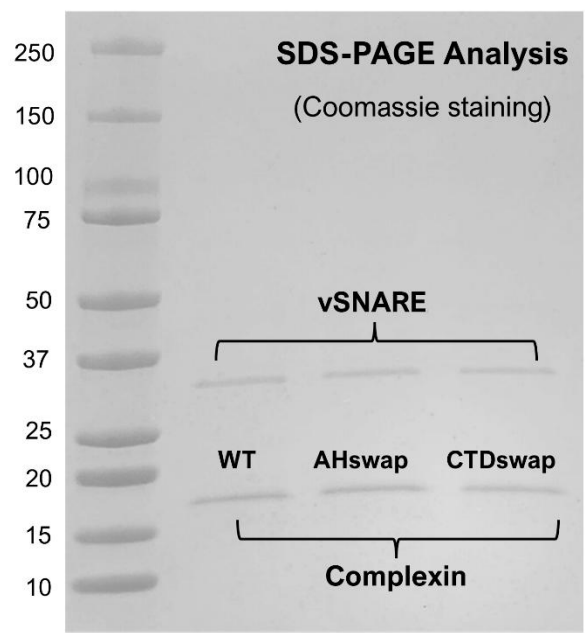

**Supplementary Figure 3.** Domain swapping does not alter CPX interaction with SNARE complexes. Binding was assessed using pull-down assays with t-SNARE complexes immobilized on agarose beads as ‘bait’ and v-SNARE together with CPX as ‘prey’. Bound v-SNARE/CPX proteins were eluted and analyzed by SDS-PAGE. Coomassie staining revealed comparable amounts of v-SNARE and CPX bound to t-SNARE beads for WT, AH<sup>swap</sup>, and CTD<sup>swap</sup> variants. A representative gel image for worm SNAREs and CPX variants is shown.

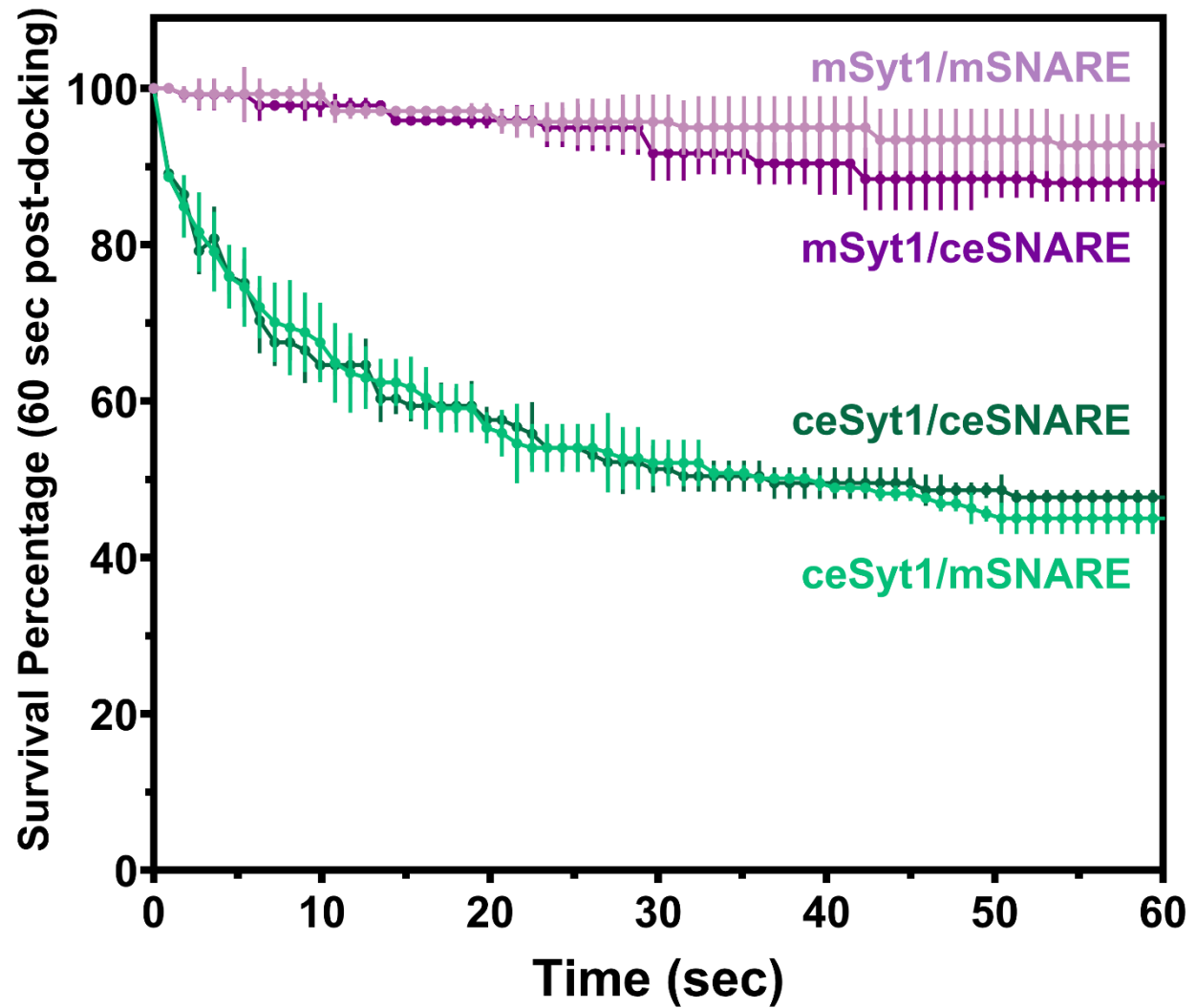

**Supplementary Figure 4.** Survival (clamping) analysis carried out under low v-SNARE conditions (~18 copies per vesicle) in the absence of CPX orthologs shows that mSyt1 (purple) is a more potent fusion clamp than ceSyt1 (green), regardless of the SNARE identity. This suggests that the difference arises from intrinsic molecular properties of the two Syt1 orthologs. Data represent mean ± SEM from three independent experiments (~180 vesicles per condition).

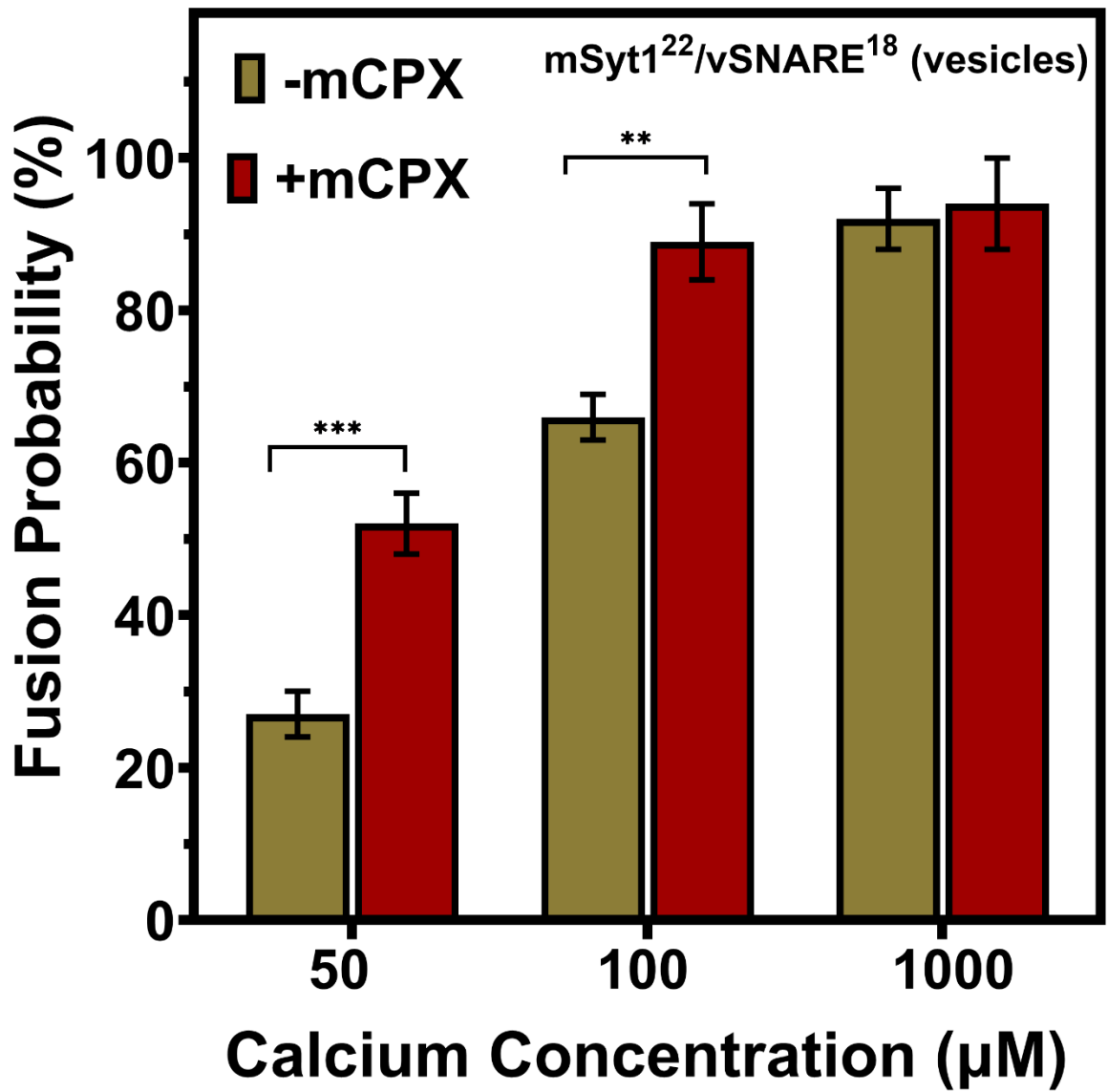

**Supplementary Figure 5.** Under low v-SNARE (~18 copies/vesicle) conditions, the Ca<sup>2+</sup>-evoked fusion probability of vesicles clamped by mSyt1 alone (green bar) was lower than in the presence of both mSyt1 and mCPX (red bar). The magnitude of this difference was inversely correlated with Ca<sup>2+</sup> concentration, pronounced at ~50 μM and negligible at 1 mM, indicating that mCPX enhances Ca<sup>2+</sup>-triggered release from mSyt1-clamped vesicles. Data represent mean ± SD from three independent experiments (~150 vesicles per condition). *Note:* Due to high spontaneous fusion under ceSyt1-only conditions, Ca<sup>2+</sup>-evoked fusion probability could not be reliably measured for the worm system.

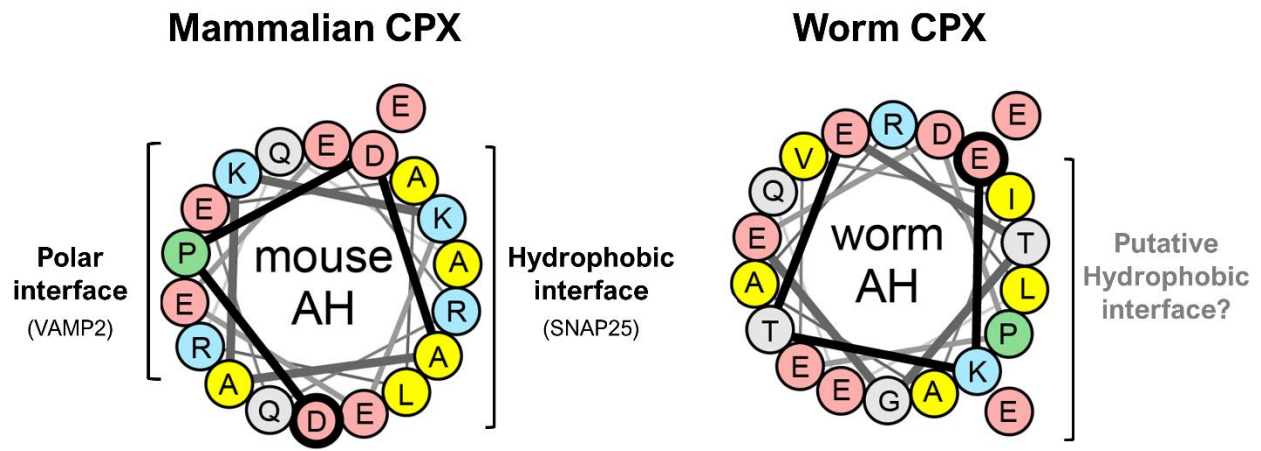

**Supplementary Figure 6.** Comparative helical wheel analysis of the accessory helix (AH) of mammalian and worm CPX, shows the AH of ceCPX contains a small hydrophobic patch analogous to the SNAP25-binding hydrophobic interface in mCPX but lacks the polar residues implicated in VAMP2 interaction. The precise mode of ceCPX AH engagement with the SNARE complex remains unknown.
